# First Molecular Verification of the Cotton Jassid (*Amrasca biguttula*) in the United States

**DOI:** 10.1101/2025.09.03.673823

**Authors:** Chaoyang Zhao, Kipling S. Balkcom

**Affiliations:** USDA-ARS, National Soil Dynamics Laboratory, Auburn, Alabama 36832, USA

**Keywords:** leafhopper, invasive insect pest, mitochondrial DNA, molecular identification, cotton agroecosystem

## Abstract

This report contains the first molecular record of the cotton jassid, *Amrasca biguttula* (Hemipter: Cicadellidae), in the United States. Nymphs across multiple instars and adult specimens were collected from a cotton (*Gossypium hirsutum*) field in Macon County, Alabama, in August 2025. While distinct paired dark spots were observed on forewings of adult specimens, this trait was present on wing pads of some nymphs but absent in others. Cytochrome oxidase I (COI) DNA barcoding confirmed the specimen identity. The United States sequence shared > 99% identity with Asian *A. biguttula* references and clustering placed the sequence within the *A. biguttula* clade with 100% posterior probability support in phylogenetic analysis. Although this pest was previously reported in 2023 from Puerto Rico based solely on morphological traits, our findings provide the first DNA-confirmed evidence of its presence in the continental United States. Given its well-documented role in damaging cotton across Asia and Africa, this report underscores the urgent need for monitoring and development of management strategies in United States cotton-growing regions.

## Main text

The cotton jassid (*Amrasca biguttula*), also known as the Indian cotton leafhopper or two-spot cotton leafhopper, is a significant insect pest that feeds on a wide range of hosts, with a primary preference for malvaceous crops such as cotton (*Gossypium spp.*) and okra (*Abelmoschus esculentus*) (Kamble and Sathe, 2015). Both nymphs and adults feed on the undersides of leaves, extracting plant sap and injecting toxic saliva. This feeding behavior induces “hopperburn”, characterized by yellowing, necrotic spotting, leaf curling, and eventual defoliation. Severe infestations can result in yield losses exceeding 60% in cotton and 50% in okra (Ahmad et al., 1986, Devi et al., 2018).

Native to Asia, this species was first reported in the Western Hemisphere in 2023, based on morphological identification of specimens collected in Puerto Rico (Cabrera-Asencio et al., 2023). In 2024 and 2025, extension bulletins and pest alerts reported the presence of *A. biguttula* in Florida, Georgia, South Carolina, and Alabama, warning growers and researchers of the potential threat. However, these early reports were based solely on morphology and the status of the pest in the United States was not previously confirmed with molecular techniques.

In August 2025, suspected “hopperburn” symptoms, including necrosis, discoloration, and leaf curling (Fig. 1A), were observed in a cotton (*Gossypium hirsutum*) field at the E.V. Smith Research Center, Auburn University, Macon County, Alabama (32.420922^o^N, 85.887618^o^W). Multiple life stages of leafhoppers were collected from the affected field (Fig. 1B-D). Adult specimens exhibited paired dark spots at the tips of their forewings (Fig. 1D), although this trait was inconsistently present in nymphal wing pads (Fig. 1B, C). Notably, the pair of small black spots typically found preapically on the crown of the head were absent in our specimens. Nymph body length varied with instar, while adults measured approximately 3 mm in length. Although these features supported a tentative identification, the morphological similarity of *A. biguttula* to related jassids necessitated molecular confirmation.

**Fig. 1.**
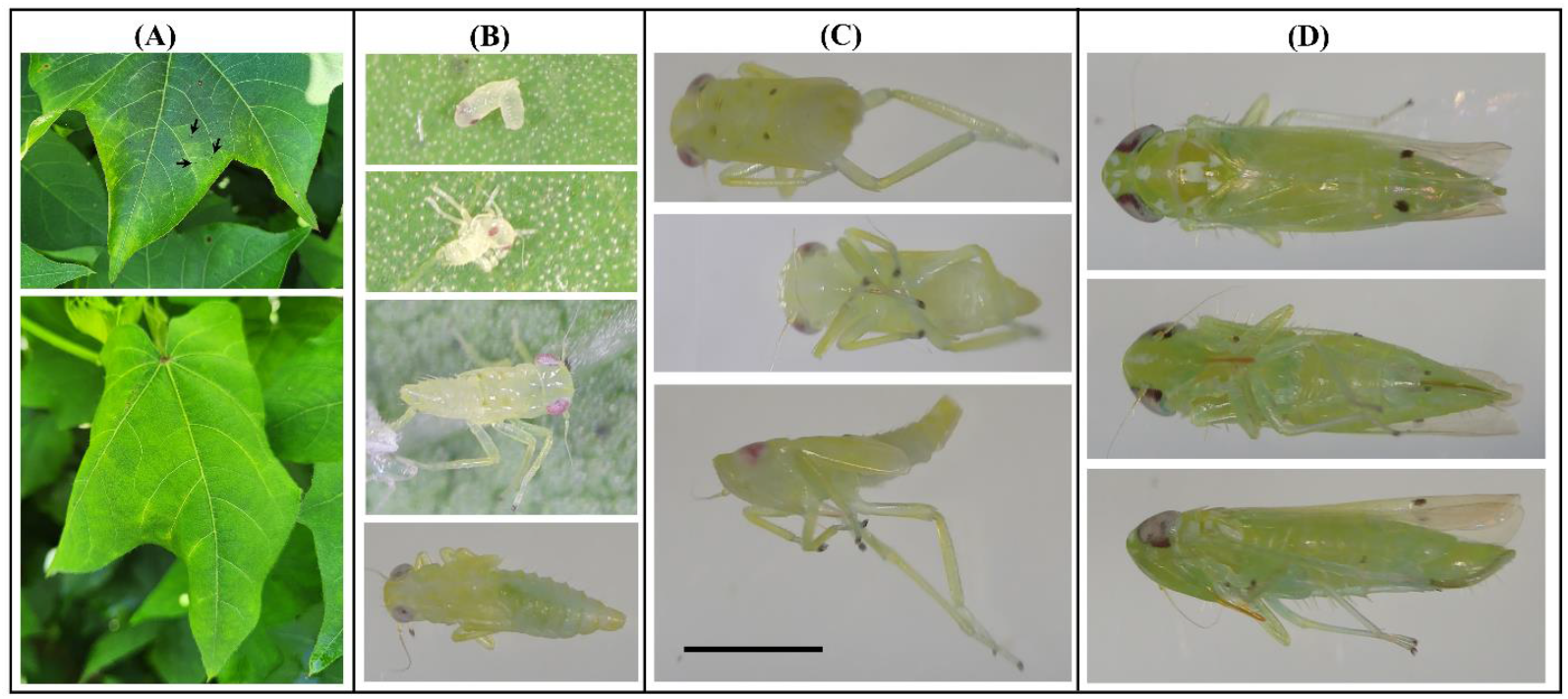
Morphology of *Amrasca biguttula* and associated cotton leaf damage. (A) Cotton leaves showing hopperburn symptoms: discoloration (top) and curling (bottom), with necrotic spotting indicated by arrows. (B) Nymphal stages: newly hatched (top), first instar (second from top), second or later instar (second from bottom), and third or later instar without wing-pad spots (bottom). (C) Later nymphal instar with wing-pad spots in dorsal (top), ventral (middle), and lateral (bottom) views. (D) Adult with forewing spots in dorsal (top), ventral (middle), and lateral (bottom) views. All insect images (B, C, D) are scaled to the reference bar (1 mm) in panel C.

Genomic DNA was separately extracted from two nymphs — one with wing-pad pigmentation (Fig. 1C) and one without (Fig. 1B, bottom) — and from one adult (Fig. 1D), using the DNeasy Blood & Tissue Kit (Qiagen, Hilden, Germany). A 658-bp barcoding fragment of the mitochondrial cytochrome oxidase subunit I (COI) gene was amplified with primers LCO1490 and HCO2198 (Folmer et al., 1994). Amplicons were cloned, and sequencing of plasmids revealed that all three individuals shared identical COI barcoding sequences, confirming they belong to the same species. The sequence has been deposited in GenBank under the accession number PX247763.

A BLAST search against the GenBank core nucleotide database returned *A. biguttula* COI sequences as the top hits. Alignment showed >99% identity between the U.S. sequence and samples collected in India, China, and Pakistan (Supplemental File 1). Consistently, Bayesian phylogenetic analysis placed the U.S. sequence within the *A. biguttula* clade with 100% posterior probability support (Fig. 2). This placement distinguished it from other *Amrasca* species as well as from non-*Amrasca* taxa within the tribe Empoascini.

**Fig. 2.**
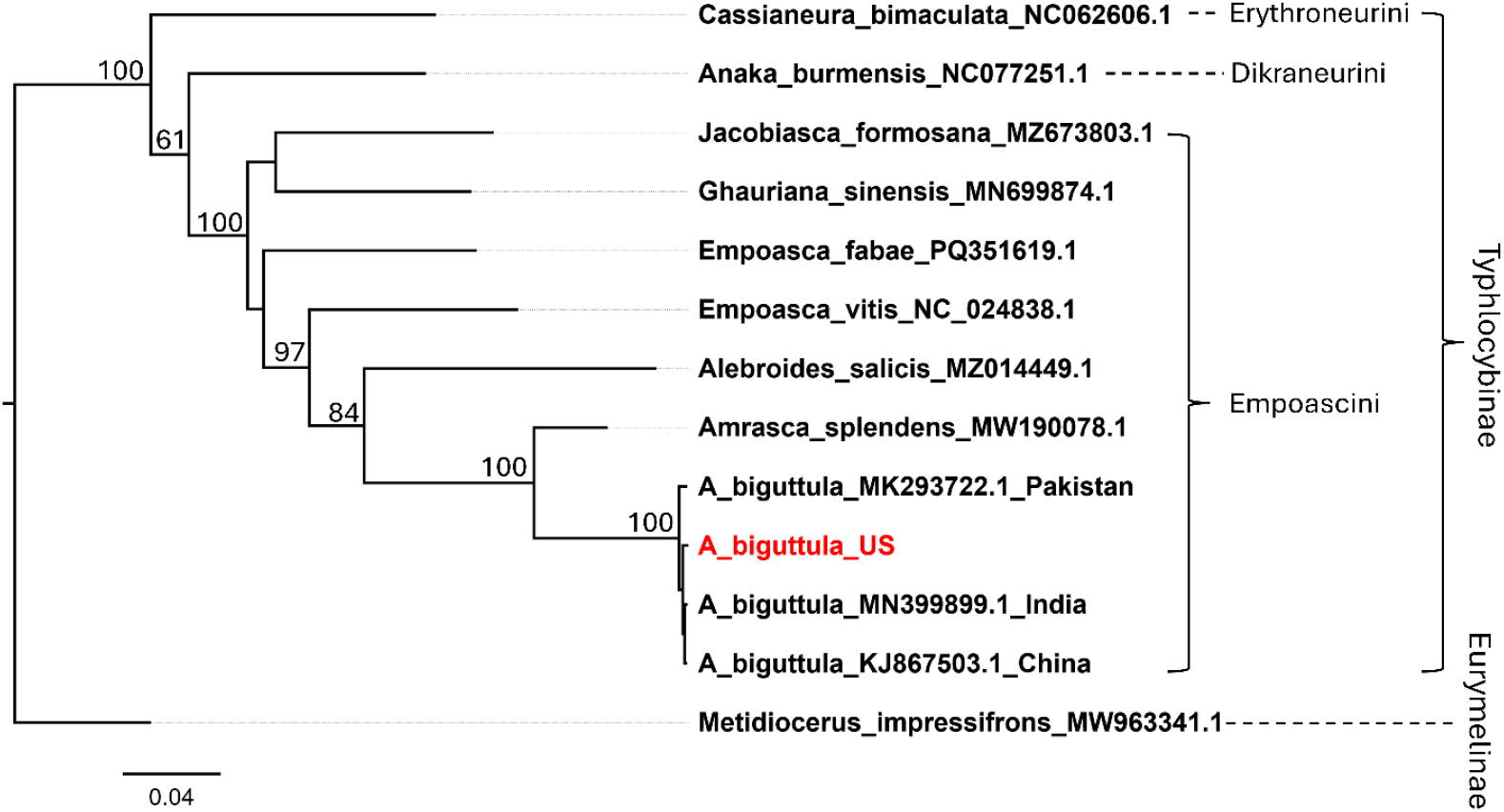
Phylogenetic placement of the U.S. *Amrasca biguttula* isolate within Asian lineages. Bayesian phylogeny of the COI barcoding region of *A. biguttula*, including the United States isolate (highlighted in red) and representative Asian sequences, alongside other species from the subfamily Typhlocybinae. The tree is rooted with *Metidiocerus impressifrons* (subfamily Eurymelinae). Taxon labels include species names followed by their corresponding GenBank accession numbers. Posterior probabilities >60% are shown at nodes. Scale bar represents substitutions per site.

The confirmation of *A. biguttula* in the United States highlights its potential to become a serious threat to cotton production. In Asia and Africa, this species is recognized as a major pest capable of causing substantial economic losses. A recent modeling study predicted that *A. biguttula* could spread and establish rapidly under different environmental conditions (Azrag et al., 2025). These findings suggest that the United States cotton-growing regions, with their warm temperatures and extensive cultivation, may be similarly vulnerable to invasion.

Despite its confirmed presence in the cotton field where *A. biguttula* was collected, we did not observe obvious yield losses at the time of sample collection during the 2025 season. This likely reflects the early stage of invasion in this location: the pest was detected after cotton had fruited, and the population had not yet propagated to high densities. Since the collection time, an insecticide was sprayed to mitigate potential yield damage. While current yield impacts appeared minimal, continued surveillance will be necessary to assess its potential to cause economic losses. Populations may increase in future seasons, which will require confirmation through accurate species identification.

Identifying *A. biguttula* adults is relatively easy due to the presence of characteristic forewing spots. However, this feature is inconsistently expressed in nymphs, which are more commonly collected during field scouting because of their limited mobility. Furthermore, cotton and other host plants may be co-infested by *A. biguttula* together with morphologically similar jassid species, making morphological diagnosis of nymphs unreliable. In such cases, molecular diagnostics offer a dependable approach of distinguishing species and accurately surveying mixed infestations to implement mitigation strategies sooner.

Molecular confirmation of *A. biguttula* in this study, based on samples collected in Alabama, provides the first molecularly validated record for the United States and serves as a valuable reference for regulatory agencies, researchers, and extension specialists. Although extension reports indicate that the pest is present in several southern states, the extent of its establishment and impact in United States cotton systems remains uncertain. Given its broad host range (Kamble and Sathe, 2015), coordinated surveillance, along with studies on management tactics such as cultural practices, biological control, and insecticide efficacy, will be essential for developing effective integrated pest management (IPM) strategies.

## Notes

### Competing Interest Statement

The authors have declared no competing interest.

