## Supplemental File 1 for "First Molecular Verification of the Cotton Jassid (*Amrasca biguttula*) in the United States"

**Supplemental File 1.** MAFFT alignment of the COI barcoding region of four *Amrasca biguttula* isoates: United States (US, this study), India (GenBank accession: MN399899.1), China (GenBank accession: KJ867503.1), and Pakistan (GenBank accession: MK293722.1).

```
US          1 AACTATATATTTTATTTTTGGGATCTGGTCTGGAATAGTAGGAATAATATTAAGTATAAT
India       1 AACTATATATTTTATTTTTGGGATCTGGTCTGGAATAGTAGGAATAATATTAAGTATAAT
China       1 AACTATATATTTTATTTTTGGGATCTGGTCTGGAATAGTAGGAATAATATTAAGTATAAT
Pakistan    1 AACTATATATTTTATTTTTGGGATCTGGTCTGGAATAGTAGGAATAATATTAAGTATAAT

US         61 CATTCGCATTGAATTAGGTCAGTTGGGTTGTTTTTTAATAAATGATCAAATATATAATGT
India      61 CATTCGCATTGAATTAGGTCAGTTGGGTTGTTTTTTAATAAATGATCAAATATATAATGT
China      61 CATTCGCATTGAATTAGGTCAGTTGGGTTGTTTTTTAATAAATGATCAAATATATAATGT
Pakistan   61 CATTCGCATTGAATTAGGTCAGTTGGGTTGTTTTTTAATAAATGATCAAATATATAATGT

US        121 TATTGTTACTTCTCATGCTTTTATTATAATTTTTTTTATAGTTATACCTATTATAATTGG
India     121 TATTGTTACTTCTCATGCTTTTATTATAATTTTTTTTATAGTTATACCTATTATAATTGG
China     121 TATTGTTACTTCTCATGCTTTTATTATAATTTTTTTTATAGTTATACCTATTATAATTGG
Pakistan  121 TATTGTTACTTCTCATGCTTTTATTATAATTTTTTTTATAGTTATACCTATTATAATTGG

US        181 TGGTTTTGGAAATTGACTTCTACCTTTAATAATTGGTGCTCCAGATATAGCTTTTCCCCG
India     181 TGGTTTTGGAAATTGACTTCTACCTTTAATAATTGGTGCTCCAGATATAGCTTTTCCCCG
China     181 TGGTTTTGGAAATTGACTTCTACCTTTAATAATTGGTGCTCCAGATATAGCTTTTCCCCG
Pakistan  181 TGGTTTTGGAAATTGACTTCTACCTTTAATAATTGGTGCTCCAGATATAGCTTTTCCCCG

US        241 ATTAAATAATATAAGATTTTGACTTTTATTACCATCTCTCTCACTTCTTATAATTAGATC
India     241 ATTAAATAATATAAGATTTTGACTTTTATTACCATCTCTCTCACTTCTTATAATTAGATC
China     241 ATTAAATAATATAAGATTTTGACTTTTATTACCATCTCTCTCACTTCTTATAATTAGATC
Pakistan  241 ATTAAATAATATAAGATTTTGACTTTTATTACCATCTCTCTCACTTCTTATAATTAGATC

US        301 ATTTGTTGAATTAGGGGCTGGTACAGGGTGAACTGTTTACCCCCCATTATCTTCTAATAT
India     301 ATTTGTTGAATTAGGGGCTGGTACAGGGTGAACTGTTTACCCCCCATTATCTTCTAATAT
China     301 ATTTGTTGAATTAGGGGCTGGTACAGGGTGAACTGTTTACCCCCCATTATCTTCTAATAT
Pakistan  301 ATTTGTTGAATTAGGGGCTGGTACAGGGTGAACTGTTTACCCCCCATTATCTTCTAATAT

US        361 TGCTCATGGAGGGGCAAGTGTAGATTTGGCTATTTTTTCCCTTCATTTAGCTGGGGTATC
India     361 TGCTCATGGAGGGGCAAGTGTAGATTTGGCTATTTTTTCCCTTCATTTAGCTGGGGTATC
China     361 TGCTCATGGAGGGGCAAGTGTAGATTTGGCTATTTTTTCCCTTCATTTAGCTGGGGTATC
Pakistan  361 TGCTCATGGAGGGGCAAGTGTAGATTTGGCTATTTTTTCCCTTCATTTAGCTGGGGTATC

US        421 TTCTATTTTAGGGGCAGTTAATTTTATTACTACTGTAATTAATATACGATGTTCTGGGTT
India     421 TTCTATTTTAGGGGCAGTTAATTTTATTACTACTGTAATTAATATACGATGTTCTGGGTT
China     421 TTCTATTTTAGGGGCAGTTAATTTTATTACTACTGTAATTAATATACGATGTTCTGGGTT
Pakistan  421 TTCTATTTTAGGGGCAGTTAATTTTATTACTACTGTAATTAATATACGATGTTCTGGGTT

US        481 AAGTTTTGATAAAATCCCTTTATTTGTGTGATCAGTTGTTATTACAGCCTTTTTATTATT
India     481 AAGTTTTGATAAAATCCCTTTATTTGTGTGATCAGTTGTTATTACAGCCTTTTTATTATT
China     481 AAGTTTTGATAAAATCCCTTTATTTGTGTGATCAGTTGTTATTACAGCCTTTTTATTATT
Pakistan  481 AAGTTTTGATAAAATCCCTTTATTTGTGTGATCAGTTGTTATTACAGCCTTTTTATTATT

US        541 ATTATCTCTCCCTGTTTTAGCAGGTGCTATTACTATGTTATTAACTGATCGAAATTTGAA
India     541 ATTATCTCTCCCTGTTTTAGCAGGTGCTATTACTATGTTATTAACTGATCGAAATTTGAA
China     541 ATTATCTCTCCCTGTTTTAGCAGGTGCTATTACTATGTTATTAACTGATCGAAATTTGAA
Pakistan  541 ATTATCTCTCCCTGTTTTAGCAGGTGCTATTACTATGTTATTAACTGATCGAAATTTGAA

US        601 TACTTCATTTTTTGATCCCTCAGGTGGAGGTGATCTAATTTTATATCAACATTTATTT
India     601 TACTTCATTTTTTGATCCCTCAGGTGGAGGTGATCCAATTTTATATCAACATTTATTT
China     601 TACTTCATTTTTTGATCCCTCAGGTGGAGGTGATCCAATTTTATATCAACATTTATTT
Pakistan  601 TACTTCATTTTTTGATCCCTCACGTGGAGGTGATCCAATTTTATATCATCATTTATTT
```
